# Integrated Clinical and Proteomic Precision Subgrouping for Severe Dengue Endotype Signature

**DOI:** 10.64898/2026.08.13.744720

**Authors:** Trupti Satish Kadni, Anoop T Ambikan, Iva Filipovic, Muralidhar Varma, Devasmita Dutta, Chiranjay Mukhopadhyay, Soham Gupta, Piya Paul Mudgal, Ujjwal Neogi

**Author notes:** Joint Correspondence. Equal Contribution (Trupti Satish Kadni, Anoop T Ambikan).

## Abstract

**Background:** Severe dengue remains difficult to predict because patients with different clinical trajectories may present with overlapping features, and conventional severity classifications may not fully capture underlying biological heterogeneity. In this study, we applied an integrated clinical and proteomic endotyping approach to dissect dengue disease heterogeneity and identify molecular signatures associated with severity.

**Methods:** Plasma proteomic profiles were analyzed together with detailed clinical, biochemical, hematological, coagulation, and immunological parameters from healthy controls and dengue patients classified according to WHO 2009 severity criteria. High-throughput proteomic analysis, unsupervised clustering, pathway enrichment, and machine-learning–based classification were used to identify dengue endotypes and define molecular features associated with predicted severe disease.

**Results:** Increasing dengue severity was associated with progressive abnormalities in liver function, coagulation parameters, hematological indices, and inflammatory mediators, including IL-6, IL-15, HGF, and MUC-16. However, proteomic profiling revealed substantial overlap across conventional severity categories, indicating that clinical classification alone does not fully resolve dengue host-response heterogeneity. Integrated clinical-proteomic clustering identified distinct dengue endotypes, including a predicted severe endotype enriched for inflammatory, antiviral, and cytotoxic lymphocyte-associated pathways. This high-risk endotype was characterized by elevated IL-15, IFN-γ, and granzymes, consistent with coordinated activation of cytotoxic lymphocyte-associated antiviral responses. Machine-learning analysis further showed that proteomic features were strong discriminators of this endotype, supporting their potential utility as biomarkers of severe host-response states.

**Conclusion:** Integrated clinical-proteomic endotyping provides molecular resolution beyond conventional severity grading and identifies immune pathways associated with severe dengue. This framework may improve biological understanding of dengue progression and support future risk stratification and biomarker development.

## Introduction

Dengue virus infection represents a major and growing global public health challenge, with an estimated 100–400 million infections annually and a rapidly expanding geographic footprint.^1^ Clinically, dengue is strikingly heterogeneous, ranging from a self-limiting febrile illness to severe dengue characterized by plasma leakage, hemorrhage, shock, and multi-organ involvement.^2^ The WHO clinical classification provides a framework for case management,^3^ but early prediction of disease progression and identification of patients at risk of severe outcomes remain limited, highlighting the need for improved biological stratification of dengue disease.^4^

Current stratification relies on clinical features and routine laboratory parameters such as platelet counts, hematocrit, liver enzymes, and coagulation markers.^5,6^ These measures largely capture downstream physiological consequences of infection but provide limited insight into the upstream molecular and immunological mechanisms driving disease progression. The pathogenesis of severe dengue is driven by complex and dynamic immune responses involving both innate and adaptive immunity. Dysregulated cytokine production, endothelial dysfunction, coagulation abnormalities, and immune-mediated tissue injury have all been implicated in severe disease manifestations. Moreover, single or small panels of biomarkers have shown limited reproducibility and predictive value across cohorts, underscoring the need for unbiased, system-level approaches to resolve dengue heterogeneity.^7^

System-level proteomics offers a powerful strategy to interrogate global immune and inflammatory responses at the protein level, enabling the identification of pathway-level alterations and disease-associated molecular signatures that are not captured by conventional assays.^8-10^ When combined with integrative analytical frameworks, such as unsupervised clustering and machine-learning–based feature selection, proteomics can facilitate the identification of biologically meaningful disease endotypes, distinct subgroups of patients defined by shared molecular mechanisms rather than clinical presentation alone.^10,11^ Endotyping has provided important insights into several immune-mediated and infectious diseases;^8,10,12^ however, biologically informed endotypes of dengue, particularly those linked to severe disease, remain poorly characterized.

In this study, we applied an integrated clinical and proteomic endotyping approach to dissect dengue disease heterogeneity and identify molecular signatures associated with severity. By combining detailed clinical and biochemical profiling with high-throughput proteomics, pathway enrichment, and machine-learning–based classification, we identified distinct dengue endotypes and characterized a predicted severe endotype with a unique immune and antiviral response profile. This integrative framework provides molecular resolution of dengue severity and advances our understanding of the immune mechanisms underlying severe disease and mechanism-anchored risk stratification.

## Results

### Patient Characteristics

The patient populations included in this study were 111 DENV-infected patients, and additionally, dengue-negative healthy controls (n=59). We defined dengue severity based on the WHO criteria 2009 as dengue without warning signs (DwoWS, n=40), dengue with warning signs (DwWS, n=64), and severe dengue (SD, n=7). There were no significant differences in age and sex among the severity groups. The DENV severity groups differed significantly across several laboratory and clinical parameters, including liver function [aspartate aminotransferase (AST), alanine aminotransferase (ALT)], renal markers (urea and creatinine), hematological indices [white blood cell (WBC), neutrophils, lymphocytes, hemoglobin (HB), hematocrit (HCT)], coagulation [Prothrombin Time (PT) and activated Partial Thromboplastin Time (aPTT)], pulse, and fever duration, as well as immunological markers (IgG, IgG:IgM). Additionally, severe clinical features including altered sensorium, organ involvement, hepatomegaly, and mucosal bleeding were more frequent in the SD group (all adjusted p values<0.05) (**Table 1**). The progressive alterations of liver enzymes, coagulation parameters, hematological indices, and immune markers across dengue severity groups indicate coordinated dysregulation in inflammatory, endothelial, and immunometabolic pathways.

**Table 1.** Clinical and demographic comparison of healthy control and dengue patients as per the WHO criteria.^3^.

| Parameter | HC | DwoWS | DwWS | SD | padj |
| --- | --- | --- | --- | --- | --- |
| N | 59 | 40 | 64 | 7 |  |
| Age | 31 (27- 37) | 27 (24-35) | 32 (24-37) | 36 (34-48) | 0.089 |
| Sex, Females, n (%) | 28 (47%) | 14 (35) | 15 (23) | 3 (43) | 0.6027 |
| AST, Median (Q1-Q3)* | - | 73 (43-153) | 131 (81-234) | 700(432-700) | **0.0003** |
| ALT, Median (Q1-Q3)* | - | 54 (35-99) | 85 (51-148) | 273 (154-700) | **0.0008** |
| Urea, Median (Q1-Q3)* | - | 15.5 (13-21) | 19 (15-25) | 35 (31-116) | **0.0008** |
| Neutrophil, Median (Q1-Q3)* | - | 1215 (920-2147) | 2120 (1360-3405) | 4780 (3833-7288) | **0.0009** |
| WBC, Median (Q1-Q3)* | - | 3350 (2425-4800) | 5150 (3850-7425) | 6800 (6050-10500) | **0.0009** |
| Lymphocytes, Median (Q1-Q3)* | - | 1240 (690-1840) | 1760 (1103-2648) | 995 (878-1278) | **0.042** |
| HB, Median (Q1-Q3)* | - | 14 (12-16) | 14 (13-15) | 10 (8-11) | **0.010** |
| HCT, Median (Q1-Q3)* | - | 40 (37-46) | 42 (38-45) | 30 (25-33) | **0.010** |
| aPTT, Median (Q1-Q3)* | - | 32 (30-36) | 37 (33-41) | 41 (38-42) | **0.008** |
| Pulse, Median (Q1-Q3)* | - | 60 (54.5-69) | 66 (59-86) | 84 (74-95) | **0.021** |
| Fever Duration in days, Median (Q1-Q3)* | - | 6 (5-8) | 7 (6-8) | 8 (7-8) | **0.036** |
| IgG (PanbioUnits), Median (Q1-Q3)* | 0.4098 (0.3375-0.4098) | 4.74 (0.29-69.89) | 49.70 (3.38-81.41) | 33.47 (22.10-76.25) | **0.036** |
| IgG:IgM, Median (Q1-Q3)* | - | 0.26 (0.01-1.33) | 0.93 (0.19-1.92) | 1.56 (0.63-2.16) | **0.048** |
| Altered Sensorium, Yes, n (%) | - | 0 | 1 (1.6) | 6 (85.71) | **<0.001** |
| Severe Organ Involvement, Yes, n (%) | - | 0 | 3 (4.69) | 7 (100) | **<0.001** |
| Hepatomegaly, Yes, n (%) | - | 1 (2.5) | 13 (20.31) | 3 (42.86) | **0.035** |
| Mucosal bleeding, Yes, n (%) | - | 0 | 4 (6.25) | 2 (28.57) | **0.039** |

\* AST: Aspartate Aminotransferase, international Units/ liter (IU/L); ALT: Alanine Aminotransferase (IU/L); WBC: White Blood Count; aPTT: Activated Partial Thromboplastin Time; HB: Hemoglobin; HCT: Hematocrit

### Epithelial damage and cytotoxic lymphocyte-associated inflammatory signature in severe dengue

The multi-organ and immune abnormalities (**Table 1**) observed across dengue severity groups necessitate high-throughput proteomics to resolve the underlying protein networks driving disease severity. To identify the systemic proteomics profile, we performed the Olink Target 96 Immuno-Oncology Panel (Olink AB, Sweden) that simultaneously detects 92 soluble and inflammatory proteins including chemokines, cytokines, interleukins, growth factors, as well as cytotoxic effector molecules such as granzymes. In the comparison between DwWS and DwoWS, 13 proteins were significantly elevated and three were significantly reduced in DwWS (adjusted p<0.05). Similarly, comparisons of SD with DwoWS and DwWS identified 22 and 17 proteins, respectively, that were significantly elevated in SD (**Fig. 1A-B** and **Table S1**). Four proteins were common to all three comparisons and significantly different across groups (**Fig. 1B**). The significant elevation of the pro-inflammatory cytokines, interleukins IL-6 and IL-15, indicates a heightened systemic inflammatory response characterized by excessive secretion of these cytokines that can lead to severe tissue damage and organ dysfunction (**Fig. 1C**). Comparisons between primary (n=70) and secondary dengue (n=41) infection showed widespread differences relative to healthy controls (**Fig. 1D-E**). While comparing the primary and secondary dengue 10 proteins were significantly higher abundance in secondary infections including Angiopoietin-2 (ANGPT2), Carbonic Anhydrase IX (CAIX), Adhesion G Protein-Coupled Receptor G1 (ADGRG1), Hepatocyte Growth Factor (HGF), Monocyte Chemotactic Protein 4 (MCP-4), Programmed Cell Death protein 1 (PDCD1), Decorin (DCN), CD4, CD5 and CD27 (**Table S2**). Given the absence of a strong global separation in proteomic profiles between DwWS and DwoWS, we quantified sample–sample dissimilarity across proteomic features using Gower distance. This analysis revealed that SD samples showed substantial proteomic similarity with a subset of DwWS, suggesting a shared host-response state across clinical severity categories. In contrast, DwWS and DwoWS samples exhibited a more homogeneous proteomic composition, reflected by lower pairwise Gower distances (**Fig. 1F**). Concurrent increases in hepatocyte growth factor (HGF) suggest engagement of regenerative or tissue-protective pathways alongside inflammation while maintaining effective antiviral responses. These findings also suggest that clinically defined dengue severity groups capture only part of the underlying proteomic heterogeneity, with substantial overlap across disease categories.

**Figure 1.**
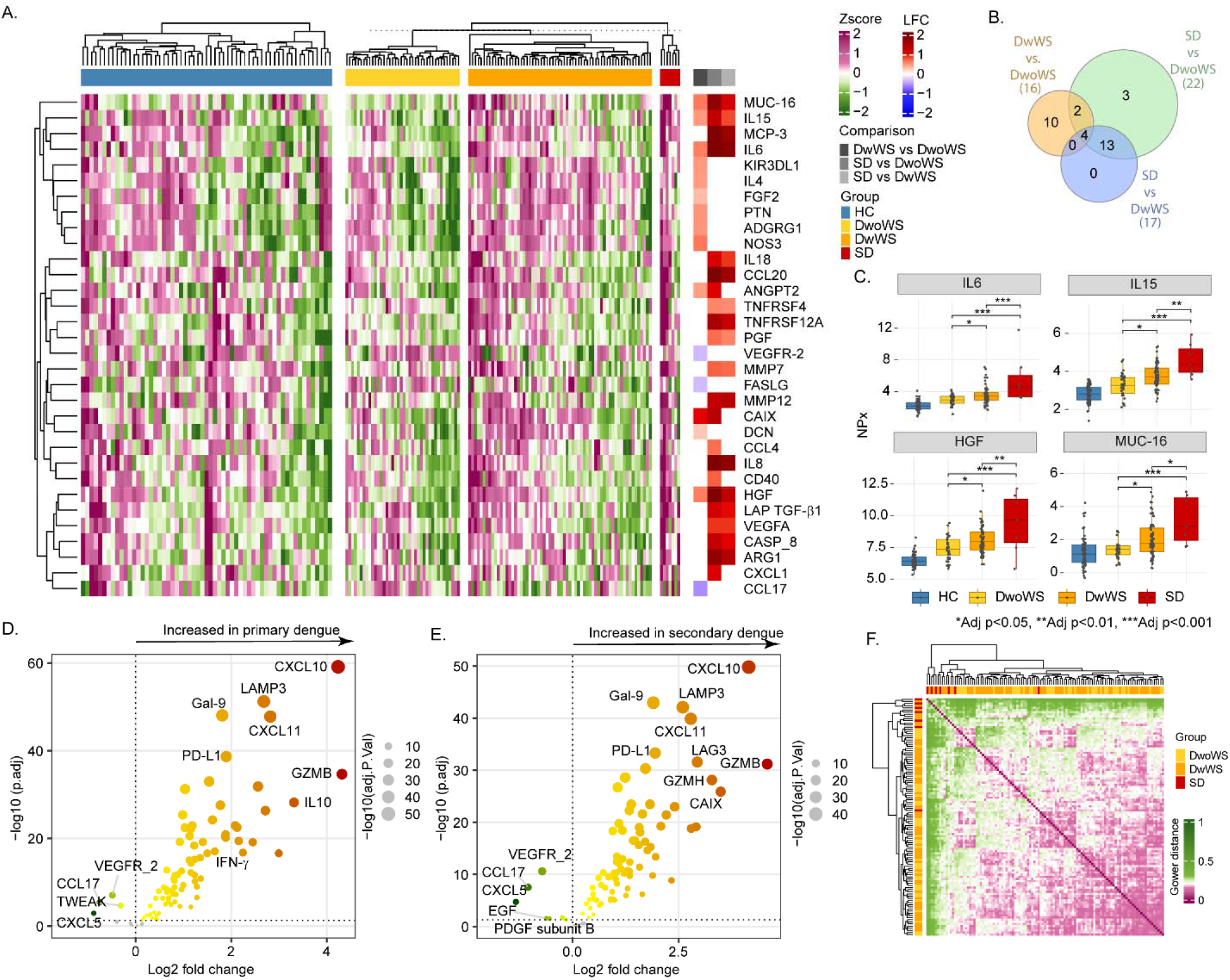
Differential protein abundance in different severity groups with DENV infection. (A) Heatmap of z-score–normalized expression for selected cytokines, chemokines, growth factors, and epithelial/inflammatory markers across all samples. Columns represent individual samples and rows represent proteins. Unsupervised hierarchical clustering was applied to both proteins and samples. The top annotation bar indicates clinical group: healthy controls (HC), dengue without warning signs (DwoWS), dengue with warning signs (DwWS), and severe dengue (SD). The sidebar summarizes the average log2fold change (LFC) per protein for the indicated comparisons. (B) Venn diagram of significantly differentially abundant proteins (adjusted *p*<0.05) shared among the pairwise comparisons: DwWS vs. DwoWS, SD vs. DwoWS, and SD vs. DwWS. Numbers indicate the count of unique and shared proteins between comparisons. (C) Boxplots of normalized protein expression (NPX) for IL-15, MUC-16, HGF, and IL-6 across clinical groups (HC, DwWS, DwoWS, SD). Boxes indicate the interquartile range with the median shown as a horizontal line; whiskers represent the full data range, and points denote individual samples. Statistical significance was assessed using multiple-testing– corrected pairwise comparisons; *Adj p* < 0.05 (\**), < 0*.*01 (**), and < 0*.*001 (\*\*\**). (D) Volcano plots comparing primary and secondary dengue infection, each relative to HC. (E) Sample-to-sample Gower distances quantifying overall dissimilarity across mixed proteomic features, where lower values indicate greater similarity.

### Precision subgroups of DENV severity

The traditional clinical severity classifications did not fully resolve the underlying molecular heterogeneity of dengue infection. We therefore sought to define molecular endotypes (subgroups of DENV patients), conceptualized here as circulating host-response subgroups, characterized by integrated clinical and biomarker profiles. The aim was to capture disease heterogeneity and identify distinct host-response patterns that are not adequately reflected by conventional clinical classifications. To define molecular endotypes, proteomic profiles were integrated with clinical and biochemical parameters. Unsupervised clustering of the combined clinical, biochemical and proteomic dataset identified three distinct endotypes, as this gave the highest average silhouette width (**Fig. 2A**). We defined them as Endotype 1, 2, and 3, where Endotype 2 comprised a small number of samples (n=4; two SD and two DwWS) and were therefore not subjected to extensive downstream comparative analyses (**Fig. 2B**). While Endotype 1 has 47.2% (17/36) of DwWS, Endotype 3 has 63.4% (45/71) of DwWS samples (**Fig. 2C**). Notably, samples from multiple clinical severity categories were represented within each endotype, indicating that the integrated molecular classification captures biologically coherent host response states that are not fully resolved by conventional clinical severity grading. Comparative analysis between Endotype 1 and Endotype 3 revealed significant differences in platelet counts; coagulation parameters (aPTT, PT); immunological markers (IgG, IgG:IgM); vital signs (pulse); liver function tests (bilirubin, AST, ALT, ALP); hematological indices (HB, HCT); and renal function markers (urea, creatinine) (adjusted p<0.05) (**Fig. 2D** and **Table 2**). The clustering identified a subgroup of individuals (Endotype 3) enriched for clinical and laboratory features associated with increased dengue severity. The proteomic data further identified broadly increased abundance of multiple immune and inflammatory proteins in Endotype 3. Together, the integration of clinical, biochemical and proteomic data highlights coordinated, endotype-specific alterations across hepatic, hematological, coagulation, and immune pathways that align with increasing dengue severity.

**Table 2.** Clinical and demographic comparison of Endotype 1 and Endotype 3.

| Parameter | Endotype 1 | Endotype 3 | Adjusted p value |
| --- | --- | --- | --- |
| Age in years, Median (Q1-Q3) | 31 (23-36) | 31 (24-37) | 0.678 |
| Sex, Female, n (%) | 15 (42) | 15 (21) | 0.163 |
| Fever Duration in days, Median (Q1-Q3) | 7 (6-8) | 6 (6-8) | 0.984 |
| PLT, Median (Q1-Q3)* | 90000 (60500-125250) | 44000 (19000-82500) | 0.0007 |
| aPTT, Median (Q1-Q3) | 32.20 (29.10-36.10) | 36.70 (33.08-41.15) | 0.0063 |
| IgG:IgM, Median (Q1-Q3) | 0.12 (0.01-0.70) | 1.06 (0.30-2.26) | 0.0063 |
| Pulse, Median (Q1-Q3) | 60 (53-70) | 70 (60-87) | 0.0089 |
| Bilirubin, Median (Q1-Q3) | 0.47 (0.33-0.86) | 0.77 (0.55-1.19) | 0.0114 |
| AST, Median (Q1-Q3) | 82 (42-145) | 140 (73-247) | 0.0137 |
| HCT, Median (Q1-Q3) | 38.95 (36.18-42.68) | 42.20 (38.30-47.05) | 0.0209 |
| IgG PanbioUnits, Median (Q1-Q3) | 1.35 (0.28-50.30) | 60.52 (3.93-77.51) | 0.0209 |
| ALT, Median (Q1-Q3) | 54.40 (39.75-75.25) | 98.00 (51.50-149.00) | 0.0214 |
| HB, Median (Q1-Q3) | 13.20 (11.80-14.48) | 14.30 (12.95-15.85) | 0.0226 |
| Urea, Median (Q1-Q3) | 15.50 (11.00-21.00) | 19.50 (15.00-27.00) | 0.0242 |
| PT, Median (Q1-Q3) | 10.80 (10.53-11.20) | 11.35 (10.73-12.08) | 0.0279 |
| Creatinine, Median (Q1-Q3) | 0.87 (0.70-0.97) | 0.96 (0.83-1.13) | 0.0325 |
| ALP, Median (Q1-Q3) | 63 (55-82) | 73 (58-123) | 0.0331 |

\*PLT: Platelets, cells/uL, AST: Aspartate Aminotransferase, international Units/ liter (IU/L); ALT: Alanine Aminotransferase (IU/L); WBC: White Blood Count; aPTT: Activated Partial Thromboplastin Time; PT: Prothrombin Time; HB: Hemoglobin; HCT: Hematocrit

**Figure 2.**
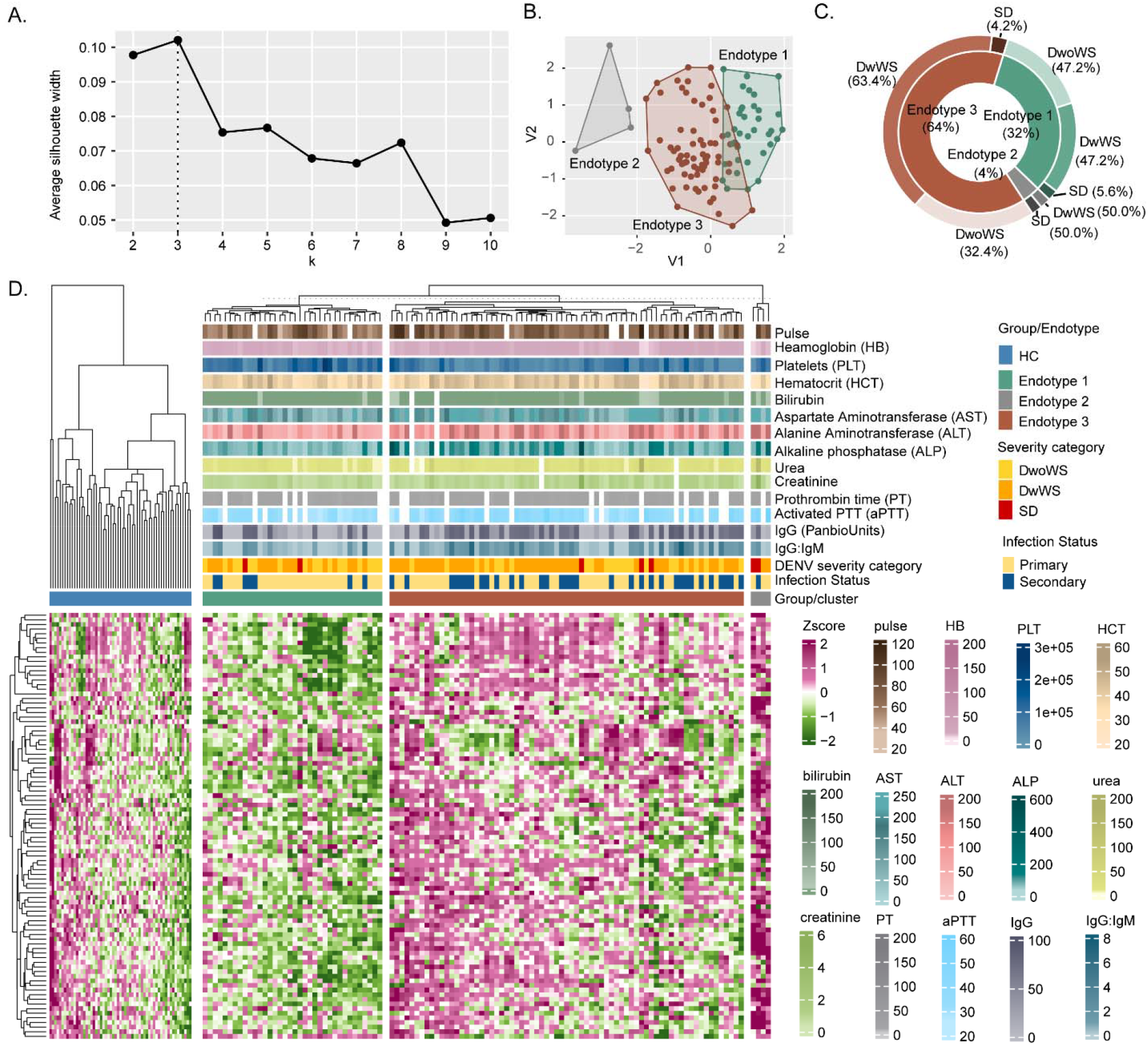
Unsupervised clustering of clinical, laboratory, and proteomic parameters identifies distinct dengue endotypes. (A) Average silhouette width analysis supporting an optimal clustering at *k* = 3. (B) Two-dimensional projection showing separation of three dengue endotypes. (C) Distribution of DENV disease severity across the identified endotypes. (D) Heatmap of z-score–scaled proteomic data, clinical, biochemical, hematological, coagulation, and immunological parameters, with hierarchical clustering of patients. Annotation bars indicate pulse, hemoglobin, platelets, hematocrit, liver and renal function markers, coagulation parameters, IgG/IgG:IgM, dengue severity category, infection status, and group/endotype assignment.

### Proteomic features associated with severe dengue endotype and risk biomarkers

To characterize proteomic features associated with the predicted severe dengue endotype, we performed the differential protein abundance analysis between Endotypes 1 and 3 after adjusting for the significantly different clinical and biochemical parameters. Among the 92 proteins tested, 60 (65%) showed significantly higher abundance in endotype 3 (adjusted p <0.05) (**Fig. 3A** and **Table S3**). The top five proteins included Granzyme B (GZMB), Granzyme H (GZMH), Interferon gamma (IFN-γ), C–C motif chemokine ligand 19 (CCL19), and Carbonic anhydrase IX (CAIX) (LFC>1.14, adjusted p<0.0001) linked to cytotoxic lymphocyte–associated immune and inflammatory signaling pathways and collectively reflect activation of cytotoxic immune programs. GZMB and GZMH are canonical cytotoxic effector molecules that are predominantly expressed by NK cells and cytotoxic T cells,^13,14^ while IFN-γ represents one of the central inflammatory cytokines, produced primarily by activated lymphocytes.^15^ Pathway enrichment analysis revealed significant activation of antiviral immune pathways, including cytokine–cytokine receptor interactions, JAK–STAT, TNF, NF-κB, innate immune sensing pathways, and NK cell–mediated cytotoxicity, alongside adaptive T cell–associated signaling (**Fig. 3B**). Random forest analysis identified coagulation parameters (aPTT and PT), platelet count, IgG, pulse rate, bilirubin, AST, and ALT as the main clinical features distinguishing Endotype 1 from Endotype 3 (**Fig. 3C**). The combined clinical model showed moderate discriminatory performance (AUC = 0.776), with aPTT, PT, IgG, platelet count, and ALT contributing most strongly (**Fig. 3D**). Proteomic analysis identified CD4, ADGRG1, IL12RB1, CASP-8, HGF, CD28, GZMH, CAIX, and NOS3 as the top discriminatory proteins (**Fig. 3E**). The combined proteomic model achieved excellent classification accuracy (AUC = 0.981), outperforming the clinical model (**Fig. 3F**). To assess whether components of this immune signature are inducible during acute dengue infection, we used a recent DENV challenge study where nine DENV-naive individuals were experimentally infected with DENV-1 strains and performed Olink Explore 3K longitudinally at 8, 10, 14, and 28 days after the infection.^16^ All the patients showed at least one systemic adverse event including leukopenia, rash and elevated liver enzymes. Granzymes and other cytokines progressively increased from day 8 post-inoculation, mirroring the abundance pattern that defines Endotype 3 (**Fig. S1**). Together, these findings support a model in which a lymphocyte activation program, characterized by co-elevation of IL-15, granzymes, and IFN-γ, defines a high-risk host-response state that spans conventional clinical severity categories.

**Figure 3.**
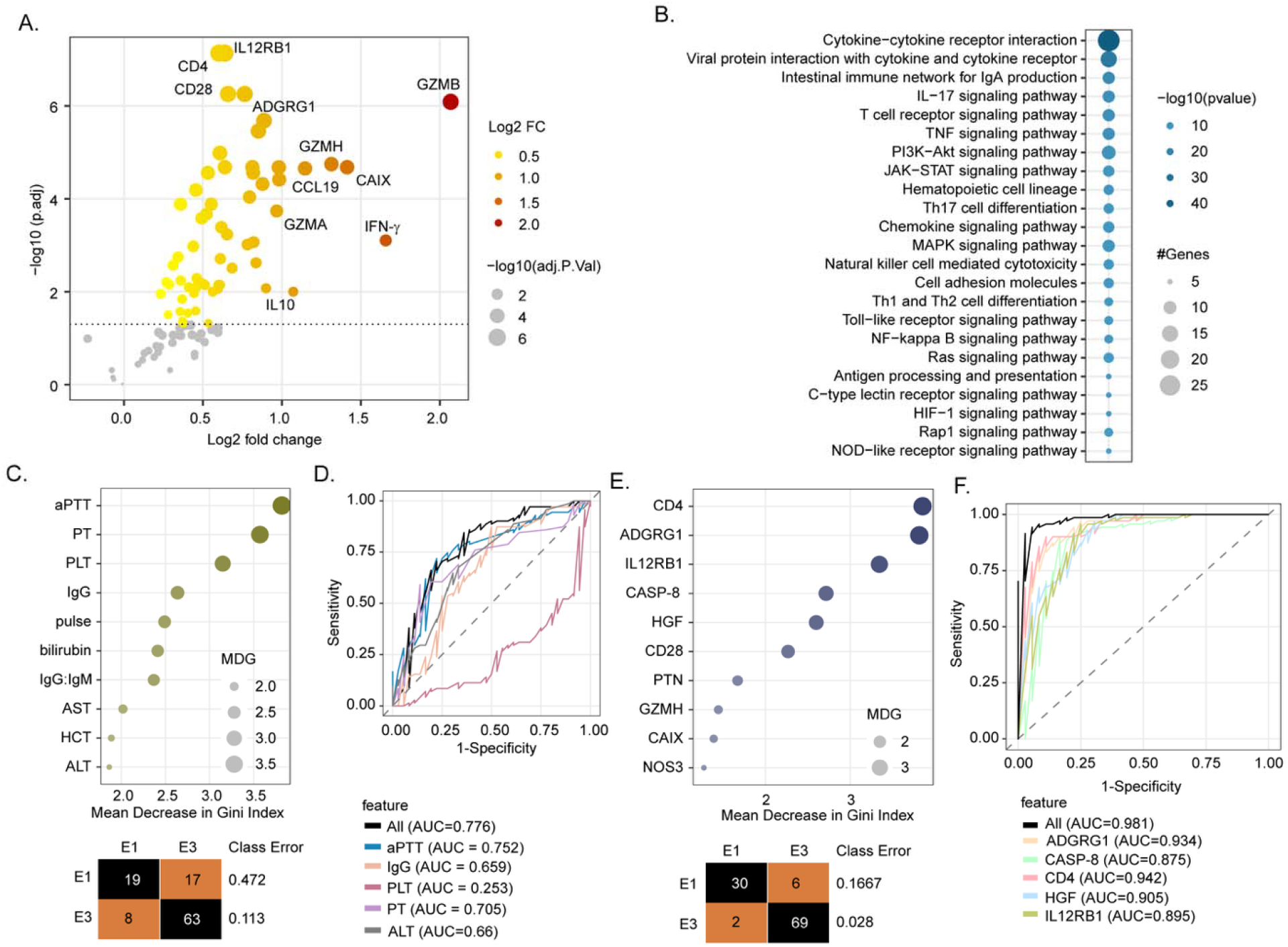
Proteomic signatures discriminate clinical endotypes in dengue infection. (A) Volcano plot of differential protein expression between Endotype 1 and Endotype 3. The x-axis shows log2fold change (LFC) and the y-axis −log10 adjusted *p*-value. Point size reflects −log10 (adjusted *p*-value), and color indicates the absolute log2fold change. Selected significantly differentially expressed proteins are annotated. (B) Pathway enrichment analysis of the differentially expressed proteins between Endotype 1 and Endotype 3. Dot size indicates the number of proteins mapped to each pathway, and color intensity represents −log10 (*p*-value), highlighting significantly enriched immune and antiviral response pathways. (C) Relative importance of clinical variables for distinguishing Endotype 1 from Endotype 3, ranked by mean decrease in Gini index from a random forest model. Coagulation parameters (aPTT, PT), platelet count, IgG level, pulse rate, and liver function markers (bilirubin, AST, ALT) emerged as the most discriminative clinical features. (D) Receiver operating characteristic (ROC) curves evaluating classification performance using clinical features. The combined clinical model achieved an AUC of 0.776, while individual features demonstrated variable discriminatory ability, with aPTT, PT, IgG, platelet count (PLT), and ALT contributing most prominently. (E) Importance ranking of selected proteomic features by mean decrease in Gini index. CD4, ADGRG1, IL12RB1, CASP-8, HGF, CD28, GZMH, CAIX, and NOS3 were the top protein features discriminating Endotype 1 from Endotype 3. (F) ROC curves for the proteomics-based classification model. The combined proteomic model achieved high discriminatory performance (AUC = 0.981), with individual proteins such as ADGRG1, CD4, IL12RB1, CASP-8, and HGF showing strong classification accuracy.

## Discussion

Severe dengue remains a major clinical challenge because early deterioration often occurs despite similar presenting features and existing severity classifications. Here, we show that integration of plasma proteomics with routine clinical and biochemical parameters resolves this challenge by uncovering molecular endotypes that more accurately reflect pathogenic host response states than conventional WHO-based severity grading. Although decreasing liver function, coagulopathy, hematological abnormalities, and immune perturbations were associated with increasing clinical severity, high-dimensional proteomic analysis revealed substantial molecular overlap across dengue severity categories, highlighting the limitations of phenotype-based classification alone. Notably, we identify a high-risk endotype defined by amplified inflammatory signaling and cytotoxic effector programs, marked by elevated IL-6, IL-15, IFN-γ, and granzymes, which together delineate a systemic immune state poised at the intersection of antiviral defense and immunopathology. Importantly, this endotype-based stratification transcends traditional clinical boundaries, with patients classified as DwWS and SD co-clustering within the same molecular state, underscoring the biological continuity of disease progression.

Our integrated clinical-proteomic analysis demonstrates that increasing dengue severity is associated with amplified systemic inflammatory signaling, most consistently reflected by elevated IL-6 and IL-15, together with markers linked to tissue stress and repair such as HGF ^17^ and MUC-16^18^. These findings are concordant with extensive clinical literature indicating that severe dengue is driven primarily by host-mediated immunopathology, rather than direct cytopathic effects of viral replication alone.^19^ IL-6 has repeatedly been shown to correlate with plasma leakage, coagulopathy, and adverse clinical outcomes in dengue patients,^20^ and experimental data support its capacity to disrupt endothelial barrier integrity and promote vascular permeability^21^. Importantly, however, IL-6 elevation is neither dengue-specific nor uniformly predictive of outcome^22,23^ reinforcing the notion that severity emerges from the magnitude, timing, and coordination of inflammatory responses, rather than the presence of any single cytokine^24,25^. In parallel with inflammatory amplification, the increase in HGF across severity strata suggests activation of counter-regulatory vascular and epithelial repair programs in response to systemic immune stress. HGF is released during endothelial and tissue injury and supports cell survival and barrier maintenance, its elevation in severe dengue likely reflects a compensatory repair response rather than inflammation alone^26^. The concurrent elevation of HGF, inflammatory cytokines, and barrier stress-associated proteins such as MUC-16 supports a model in which dengue progression involves both immune-mediated injury and an insufficient or temporally uncoupled repair response. This is consistent with the clinical observation that vascular dysfunction and plasma leakage peak during the critical phase, often as viral loads decline, highlighting the central role of host-response dynamics and barrier vulnerability in clinical deterioration.^27^ Together, elevated IL-6/IL-15–linked inflammatory signals with HGF and MUC-16 suggest that severe dengue reflects an imbalance between inflammatory injury and barrier/repair capacity, rather than uncontrolled inflammation alone.

Comparison of primary and secondary dengue patients with healthy controls revealed a broad, largely overlapping inflammatory signature, with most immune mediators significantly perturbed relative to HC. Direct comparison of secondary versus primary dengue identified only a smaller set of differences, including ANGPT2, CD27, CAIX, ADGRG1, CD5/CD4, HGF, MCP-4, PDCD1, and DCN, suggesting modest serostatus-associated shifts within a shared inflammatory response. Consistent with previous studies, these findings indicate that although secondary dengue is associated with increased risk of severe disease, disease progression is more closely linked to the magnitude, persistence, and coordination of inflammatory signaling than infection history alone^28^. The limited qualitative separation between primary and secondary dengue further supports an endotype-based model in which inflammatory burden is necessary but not sufficient for severe outcomes.

These findings should be interpreted in the context of the temporal evolution of cytokine responses in dengue. In our earlier longitudinal analysis during the early febrile phase, we identified a severity-associated cytokine signature as early as day 3 post–symptom onset, including elevations in IL-6, IL-8, IL-10, GM-CSF, IL-13, and IL-4 with reductions in IL-12 and MIP-1β in patients who progressed to severe disease.^25^ In the present study, IL-6 again emerges as a severity-linked inflammatory mediator, indicating a shared axis of inflammatory amplification that extends beyond the early febrile window. In contrast, cytokines showing early, transient polarization were less prominent in this cross-sectional analysis, likely reflecting differences in sampling timing (consistent with the fever duration at presentation in this cohort, median approx. 6-8 days) and the transition from early immune polarization to broader host-response states. Importantly, the broader proteomics profiling resolves additional mediators not assessed in the earlier panel, including IL-15 and cytotoxic inflammatory effectors, which define a high-risk dengue endotype and are consistent with later-phase immune activation associated with disease severity^29^.

Beyond conventional severity groupings, our endotype-based stratification reveals a high-risk host-response state characterized by enrichment of IL-15, granzymes (GZMB, GZMH), and IFN-γ, together with pathway enrichment for antiviral and cytotoxic immune programs. IL-15 is a central cytokine for the maintenance and activation of cytotoxic lymphocyte compartments, particularly NK cells, but also memory CD8+ T cells,^30,31^ and elevated IL-15 has been repeatedly associated with more severe dengue phenotypes and heightened cytolytic immune activity^29,32^. Granzymes, IFN-γ, and related effector proteins are expressed by several cytotoxic lymphocyte subsets, including NK cells, CD8^+^ T cells, gamma delta (γδ) T cells, MAIT cells, and the significantly enriched pathways identified in our study could be attributed to any of these. Plasma proteomic signature can therefore not be attributed to a specific immune cell lineage. However, among these subsets, NK cells have been some of the most extensively characterized in dengue infection. One study has shown elevated granzyme B, IFN-γ and CD69 in CD56^dim^ NK cells as well as higher soluble IL-15 in DwoWS compared to SD,^33^ and another one reduced granzyme B and degranulation capacity in SD.^34^ These results are not necessarily contradicting ours, as during the sustained in vivo degranulation upon onset of infection, granule release would present as elevated soluble effector molecules (e.g., granzymes) in plasma. Consistent with this, intracellular granzyme B was reported at similar expression levels across severity categories and healthy controls, despite increased degranulation capacity.^35^ Other work reported that NK cells retain their functional capacity during the acute dengue infection.^36^ Differences in sampling time are likely to contribute further, as the prior studies assessed the early febrile phase, and patients in our study presented after a median fever duration of 6-8 days. Notably, ADGRG1, used as a cytotoxicity marker for NK cells in SD,^34^ was among the strongest proteomic discriminators of Endotype 3 in our data, providing a link between circulating and cellular compartments, though this link warrants further investigation. CD4^+^ T cells also merit consideration here. A subset of CXCR5^-^PD-1^hi^ CD4^+^ T cells was recently identified to accumulate in SD and DwWS patients.^37^ The CD4 severity-discriminating signal in our data marks these cells as a further candidate source of the mediators enriched in Endotype 3. Resolving whether Endotype 3 reflects lymphocyte activation, exhaustion, or both requires deep functional and phenotyping profiling. The prominence of granzymes and IFN-γ in Endotype 3 is therefore consistent with a state of cytotoxic immune activation, rather than definitive evidence of a single causal cell type.

The finding that samples from different WHO severity categories cluster within the same endotype indicates that clinical severity groups do not map one-to-one onto underlying immune profiles. Instead, patients classified by conventional criteria can share similar circulating host-response states, reflecting overlaps rather than discrete biological categories. This interpretation is consistent with longitudinal studies demonstrating that inflammatory and interferon-associated signals change over time and that a subset of patients shows sustained cytotoxic and chemokine-related signatures preceding clinical worsening^38,39^. Supporting the plausibility of this endotype, analysis of a controlled DENV-1 human challenge cohort showed temporal induction of granzymes and inflammatory cytokines in antigen-naive individuals, indicating that elements of the observed cytotoxic signature can emerge during acute infection.

Pathway enrichment analyses further highlighted JAK-STAT and NF-κB signaling, which are central hubs integrating cytokine and interferon responses during dengue infection^40^. Both pathways are well-established mediators of antiviral defense and inflammatory amplification, and their activation has been linked to disease severity and endothelial dysfunction in dengue and other viral infections^32,41^. Importantly, IL-6 and IL-15 signal directly through JAK-STAT pathways^42^, while IFN-γ engages both JAK-STAT and NF-κB-dependent transcriptional programs,^43^ providing a mechanistic framework linking the observed cytokine milieu with downstream inflammatory and cytotoxic gene expression. At the same time, NF-κB-driven inflammation can potentiate vascular permeability and coagulation disturbances when dysregulated^44^, suggesting that sustained activation of these signaling axes may contribute to immunopathology rather than protection in high-risk dengue host-response states.

This study has several limitations. First, the severe dengue group was relatively small, which may limit statistical power and the generalizability of severity-associated proteomic signatures. Therefore, validation in larger and independent dengue cohorts is needed to confirm the robustness of the identified endotypes and candidate biomarkers. We have initiated a large multi-center study to validate the endotype based severity biomarkers (COMBAT, ClinicalTrials.gov ID NCT06751836).^45^ Second, the cross-sectional design limits our ability to determine the temporal sequence between immune activation, molecular endotype formation, and subsequent clinical deterioration. Although longitudinal sampling would provide greater insight into how these host-response states evolve over the course of infection, such sampling is challenging in real-world dengue care because patients present to hospital at variable times after symptom onset. Importantly, time from symptom onset to sampling was not significantly different across the identified endotypes, suggesting that the observed molecular subgroups were not primarily driven by sampling time. Nevertheless, future longitudinal studies are required to determine whether these endotypes precede, accompany, or predict progression to severe dengue. Finally, plasma proteomics capture circulating immune and inflammatory signatures but cannot definitively identify the cellular sources of key mediators such as IL-15, IFN-γ, and granzymes. Future studies integrating single-cell profiling, cellular immunophenotyping, or functional assays will be important to resolve the immune cell populations and mechanisms driving these high-risk host-response states.

In conclusion, our study demonstrates that severe dengue is best understood as a continuum of host-response states^46^ rather than as discrete clinical categories defined solely by conventional severity grading. By integrating plasma proteomics with routine clinical and biochemical parameters, we identify a high-risk endotype characterized by coordinated inflammatory, JAK-STAT/NF-κB, and cytotoxic immune programs, including elevated IL-6, IL-15, IFN-γ, and granzymes. The co-clustering of DwWS and SD patients within this endotype highlights the biological overlap underlying clinical progression and suggests that molecular stratification may better capture patients at risk of deterioration. Together, these findings support a model in which severe dengue reflects an imbalance between antiviral immune activation, inflammatory injury, and insufficient barrier or tissue repair responses. Endotype-based classification may therefore provide a more precise framework for risk prediction and future therapeutic targeting in dengue.

### Methodology

#### Ethical Approval and Participant Enrolment

This study was proposed and approved by Kasturba Medical College and Kasturba Hospital Institutional Ethics Committee (approval number IEC1-204-2022), Institutional Biosafety Committee, Manipal Academy of Higher Education, Clinical Trials Registry-India (CTRI/2022/10/046293) and the Swedish Ethical Review Authority (Ref. number: 2024-05758-01). Study participants-both DENV-positive patients and DENV-negative healthy controls between the age of 18-60 were enrolled to the study after obtaining their informed consent. The detailed description of the study and the intent of the sample collection was explained to the participants. With the help of a detailed questionnaire, the patient past health and known co-morbidities, current infection and ongoing symptoms, any previous hospitalization for the current infection and any history of recent vaccination including COVID-19 vaccination was collected from the patients and controls appropriately. In the case of DENV-patients, individuals tested positive for dengue with the help of serological investigations such as IgM ELISA, NS1 ELISA or both and in some cases by molecular methods such as RT-PCR, were approached for informed consent. The patient health and recovery were tracked during their course of hospitalization to note the onset of any new warning signs and/ or assess improvement in their condition. The clinical discharge summary of patients was accessed after the necessary ethical approvals through the Medical Records Department, Kasturba Hospital to ensure that all data points were captured in case the patient failed to mention any past symptoms that they may have recovered from, questionnaire interviewer bias and in case of severe patients where the family consented on behalf of the patient.

#### Sample Size

Due to the complexity of the high dimensionality involved in omics-based studies, this study relied on purposive sampling to calculate the required sample size. The patients enrolled to the study were categorized on the day of enrolment to the study as per the WHO 2009 categorization into one of three groups.

#### Patient Blood Profile Investigations

Routine hematological investigations to track thrombocytopenia were performed daily to monitor patient health and potentially adjust treatment strategies. Additional testing of biochemical panels capturing organ health and blood coagulation was risk-stratified based on disease severity and was performed based on the discretion of the treating clinician. Accordingly, patients with dengue warning signs or severe dengue underwent tests such as prothrombin time (PT), activated partial thromboplastin time (aPTT), procalcitonin etc. Relevant laboratory results were accessed and recorded through the Medical Records Department at Kasturba Medical College and Hospital, Manipal.

The list of tests are as follows: Hematology: Hemoglobin, White Blood Cells (WBC), lymphocytes, platelets, hematocrit; Biochemistry: bilirubin, Aspartate Transaminase (AST), Alanine Transaminase (ALT), Alkaline Phosphatase (ALP), urea, creatinine, Serum Lactate Dehydrogenase (LDH), Creatinine Phosphokinase (CPK); Coagulation: Prothrombin Time (PT), Activated Partial Thromboplastin Time (aPTT); Cardiac: Troponin T, N-Terminal Pro-B-Type Natriuretic Protein (NT-proBNP); Inflammation/Infection: Procalcitonin; Blood gas/metabolic: Lactate Oxidase (Arterial Blood Gas). The reference Range is given in the supplementary Table S4.

#### Patient Severity Scoring

Patient severity was assessed during patient enrolment to the cohort and until their discharge. This was done specifically to track and identify potential defervescence that is often recorded in dengue infections. The WHO Tropical Disease Research 2009 classification was used for the categorization of patients into the following three groups and their respective symptoms: 1. Dengue without warning signs (DwoWS), 2. Dengue with warning signs (DwWS), 3. Severe dengue (SD). The highest recorded score of severity was noted as the category and the patients were not moved down on recovery. In case of development of further warning signs, depending upon the type of sign, the patient was appropriately shifted up on the severity grading scale. Based on this severity grading, 40 patients from DwoWS, 64 patients from DwWS and 7 patients in SD groups were included in the study, along with 59 healthy control samples.

#### Antibody and Cytokine Analysis

The plasma samples collected from patient and control groups were used for IgM, IgG and proteomic investigations. Panbio anti dengue IgG ELISA was carried out as per kit protocol following manufacturer’s instructions. The identification of secondary cases was performed using the cut off at 22 Panbio units. Proteomics investigations were conducted using 92 inflammatory proteins was determined using proximity extension immuno-assay technology (Olink Proteomics®) with the Olink Target 96 Immuno-oncology panel as per manufacturer’s protocols.

#### Bioinformatics and statistical analysis

Differential protein abundance analysis of normalized protein expression (NPX) data was performed using the limma R package (v3.60.4)^47^. To characterize dengue disease endotypes, targeted proteomic data (n=92 variables) were integrated with clinical and laboratory data (n=48) available for a subset of patients drawn from a total cohort of 111 dengue-infected individuals. The integrated dataset comprised 48 variables in total, including 25 categorical variables and 23 continuous variables. Categorical variables captured demographic characteristics, clinical symptoms, lifestyle factors, and disease manifestations. Continuous variables included demographic measures, vital signs, hematological indices, and biochemical, coagulation, and immunological parameters. Sample similarity was assessed using the Gower distance metric to accommodate mixed data types, implemented via the daisy function in the R package cluster (v2.1.6). The resulting distance matrix was visualized as a heatmap to examine global patterns of inter-sample similarity. Unsupervised clustering was performed using agglomerative hierarchical clustering with Ward’s linkage method using the agnes function from cluster R package (v2.1.6). The optimal number of clusters was determined by evaluating average silhouette widths across a range of cluster solutions (k = 2-10), computed using the silhouette R function. Based on this criterion, the dataset was partitioned into three clusters using the cutree R function. Cluster structure was further visualized using classical multidimensional scaling implemented with cmdscale function, projecting the Gower distance matrix into two dimensions for visualization of endotype separation. Continuous variables were compared between groups using the nonparametric Mann–Whitney U test, implemented in the R package rstatix (v0.7.2). Categorical variables were assessed using Fisher’s exact test. Multiple testing correction was applied using the Benjamini–Hochberg method. To investigate underlying biological mechanisms, pathway enrichment analysis was performed using the Enrichr framework via the Python package gseapy (v1.1.5)^48,49^. KEGG pathway gene sets were used, focusing on metabolic, environmental information processing, and organismal system pathways. Pathways were considered significantly enriched at an adjusted p-value<0.05 with a minimum overlap of five proteins. To identify discriminative molecular and clinical features, random forest classification models were constructed separately for proteomic and clinical datasets using the R package randomForest (v4.7.1.2). Each model was trained using 1,000 decision trees and included all available samples and features. Predictive performance was evaluated using receiver operating characteristic (ROC) curve analysis, and model discrimination was quantified using the area under the curve (AUC). Variable importance was assessed using the Gini impurity index.

## Acknowledgements

We gratefully acknowledge Manipal Institute of Virology for providing the necessary institutional research funds and infrastructural facilities to support this work. We also thank and acknowledge the clinicians, nursing staff and the study participants of Kasturba Hospital, Manipal for their invaluable assistance and cooperation throughout the study. This study has received funding from the European Union’s HORIZON Research and Innovation Actions under grant agreement No. 101191315 (COMBAT). Views and opinions expressed are, however, those of the authors only and do not necessarily reflect those of the European Union. The European Union cannot be held responsible for them. UN acknowledges support received from the Swedish Research Council grants (2021-01756, 2021-00993, 2018-06156 and 2017-01330), Karolinska Institutet Consolidator Grant (2-117/2023). IF acknowledges support received from Swedish Research Council Starting Grant (2025-02198), Jeanssons Stiftelser (J2025-0113), and Åke Wibergs Stiftelse (M25-0374). We thank Olink™ Ab for providing the support to run the proteomics as a partnership project at Premas Bioscience Laboratory, New Delhi. Olink does not have any role in study design, interpretation or analysis of the data.

## Authors Contribution

Conceptualization of clinical study: T.S.K., M.V., C.M., P.P.M., and U.N.

Conceptualization of the bioinformatics part, A.T.A., I.F., and U.N.

Methodology: T.S.K., A.T.A., I.F., D.D., S.G., R.L.J., S.W., and U.N.

Investigation: M.V., and C.M.

Clinical data management: T.S.K., M.V., C.M., and P.P.M.

Visualization: A.T.A., I.F., and U.N.

Supervision: P.P.M., and U.N.,

Writing-original draft: T.S.K, A.T.A., S.G., I.F., and U.N.

Writing-review & editing: All authors

## Notes

### Competing Interest Statement

The authors have declared no competing interest.

