## Supplementary Fig S1 and Table S4 for "Integrated Clinical and Proteomic Precision Subgrouping for Severe Dengue Endotype Signature"

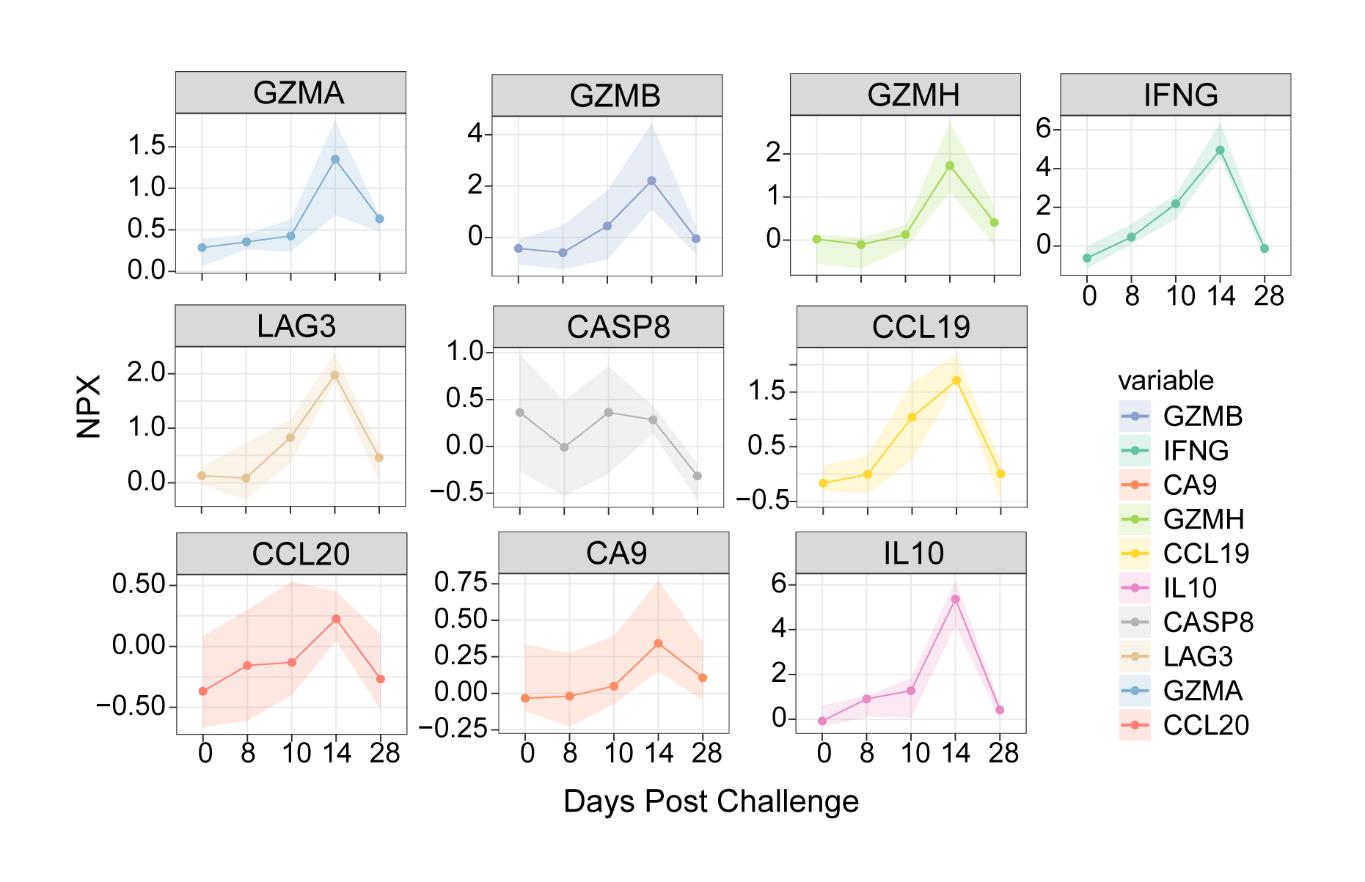

Figure S1. Temporal protein abundance of the selected proteins post-DENV challenge assay.

Table S4. Normal range of the test for Indian population based on the test used.

| Test | Unit of Measurement | Normal Range Male | Normal Range Female |
| --- | --- | --- | --- |
| Hemoglobin | g/dL | 13-17 | 12-15 |
| White Blood Cells | (value)/µL | 4000-10000 | 4000-10000 |
| Differential Lymphocyte Count | (value)/µL | Neutrophils | 2000-8000 |
|  |  | Eosinophils | 40-400 |
|  |  | Lymphocyte | 720-4400 |
| Platelets | (value)/µL | 150000-400000 | 150000-400000 |
| Hematocrit | % | 40-50 | 34-46 |
| Procalcitonin | mcg/L | Less than 0.5 | Less than 0.5 |
| Total Bilirubin | mg/dL | Up to 1.2 | Up to 1.2 |
| Aspartate Transaminase | IU/L | Up to 40 IU/L | Up to 32 IU/L |
| Alanine Transaminase | IU/L | Up to 41 IU/L | Up to 33 IU/L |
| Alkaline Phosphatase | U/L | 40-130 | 35-105 |
| Urea | mg/dL | 16.6-48.5 | 10-40 |
| Creatinine | mg/dL | 0.7-1.2 | 0.5-0.9 |
| Partial Thromboplastin Time | seconds | - | - |
| Activated Partial Thromboplastin Time | seconds | - | - |
| Serum Lactate Dehydrogenase | U/L | 135-285 | 135-214 |
| Lactate Oxidase (Arterial Blood Gas) | mg/dL | 5-14 | 5-14 |
| Creatinine Phosphokinase | U/L | 46-171 | 34-149 |
| Troponin T | ng/mL | Up to 0.020 | Up to 0.020 |
| N-Terminal Pro- B-Type Natriuretic Protein | pg/mL | 40-125 | 40-125 |
